# Structure of the cysteine-rich domain of *Plasmodium falciparum* P113 identifies the location of the RH5 binding site

**DOI:** 10.1101/2020.06.12.149534

**Authors:** Ivan Campeotto, Francis Galaway, Shahid Mehmood, Lea K. Barfod, Doris Quinkert, Vinayaka Kotraiah, Timothy W. Phares, Katherine E. Wright, Ambrosius P. Snijders, Simon J. Draper, Matthew. K. Higgins, Gavin J. Wright

## Abstract

*Plasmodium falciparum* RH5 is a secreted parasite ligand that is essential for erythrocyte invasion through direct interaction with the host erythrocyte receptor basigin. RH5 forms a tripartite complex with two other secreted parasite proteins: CyRPA and RIPR, and is tethered to the surface of the parasite through membrane-anchored P113. Antibodies against RH5, CyRPA and RIPR inhibit parasite invasion, suggesting that vaccines containing these three components have the potential to prevent blood-stage malaria. To further explore the role of the P113-RH5 interaction, we selected monoclonal antibodies against P113 that were either inhibitory or non-inhibitory for RH5 binding. Using a Fab fragment as a crystallisation chaperone, we determined the crystal structure of the RH5-binding region of P113 and showed that it is composed of two domains with structural similarities to rhamnose-binding lectins. We identified the RH5 binding site on P113 by using a combination of hydrogen-deuterium exchange mass spectrometry and site directed mutagenesis. We found that a monoclonal antibody to P113 that bound to this interface and inhibited the RH5-P113 interaction did not inhibit parasite blood-stage growth. These findings provide further structural information on the protein interactions of RH5 and will be helpful in the development of blood-stage malaria vaccines that target RH5.

**Importance:** Malaria is a deadly infectious disease primarily caused by the parasite *Plasmodium falciparum.* It remains a major global health problem and there is no highly effective vaccine. A parasite protein called RH5 is centrally involved in the invasion of host red blood cells, making it – and the other parasite proteins it interacts with – promising vaccine targets. We recently identified a protein called P113 that binds RH5 suggesting that it anchors RH5 to the parasite surface. In this paper, we use structural biology to locate and characterize the RH5 binding region on P113. These findings will be important to guide the development of new anti-malarial vaccines to ultimately prevent this disease which affects some of the poorest people on the planet.

## Introduction

Malaria is a devastating infectious disease caused by parasites from the genus *Plasmodium*, and in 2018, it was responsible for an estimated 228 million clinical cases (1). Over 99% of malaria cases are caused by *Plasmodium falciparum*, a parasite that is endemic in many tropical regions and is responsible for over 400,000 deaths each year (1). While several licenced drugs kill *Plasmodium* parasites, the requirement to treat each new infection, and the emergence of drug-resistant parasites, threaten current control methods (2). A vaccine that elicits high levels of long-lasting protection will be a valuable tool in the battle against malaria.

The symptoms of malaria occur when the parasite replicates within human blood. This is initiated when the merozoite form of *Plasmodium* recognises and invades a host erythrocyte. Invasion requires molecular interactions between parasite ligands, which are released in an ordered schedule from intracellular organelles, and receptor proteins displayed on host erythrocyte surfaces (3, 4). As erythrocyte invasion is an essential stage of the parasite life cycle, and the merozoite is directly exposed to host antibodies, invasion has long been considered a suitable target for vaccine-elicited antibodies. An important advance in targeting the blood-stage was the discovery that the parasite protein, reticulocyte-binding protein homologue 5 (RH5), makes an interaction with erythrocyte basigin which is essential and universally required by all strains of parasite for invasion (5). This interaction has been structurally characterised (6) and studies have shown that anti-RH5 antibodies can prevent erythrocyte invasion by multiple *Plasmodium falciparum* strains (7–9). Vaccination of non-human primates with RH5 protected them from challenge with a heterologous parasite strain (10) and anti-RH5 monoclonal antibodies can passively protect non-human primates (11). While human clinical trials of RH5 are underway (12), the analysis of antibodies, elicited through human vaccination, has been instructive in revealing the epitopes of protective and potentiating antibodies which should be induced by future focused vaccines (7).

RH5 does not act alone on the surface of the merozoite, but forms a tripartite complex with two other secreted parasite proteins: cysteine-rich protective antigen (CyRPA) (13–15) and RH5-interacting protein (RIPR) (16). Prior to invasion, these proteins are spatially segregated: RH5 is sequestered within the rhoptry (17) and both CyRPA and RIPR are localised to the micronemes (14). The proteins ultimately co-localise, most likely at the point of invasion, and the complex has been studied using recombinant proteins in binary protein interaction assays (18) and its architecture determined to 7Å resolution (19). Recently, a fourth interacting partner of RH5 was identified as an abundant GPI-anchored merozoite surface protein called P113 and shown to tether the RH5:CyRPA:RIPR complex to the merozoite surface (18). The interaction is conserved across the *Laverania* subgenus (20), and the core of the interaction was mapped to the N-terminal region of P113 (residues 1-197) and a 19-residue peptide from the flexible and disordered N-terminus of RH5 (residues 9-27) which does not interact with RIPR or CyRPA. Polyclonal antibodies raised against the RH5 N-terminus (residues 1-116) inhibited the interaction with P113 and also inhibited parasite growth *in vitro* (18). It was not known, however, whether antibodies that target P113 and prevent it from binding to RH5, would also prevent erythrocyte invasion. We therefore raised monoclonal antibodies against P113 which are inhibitory and non-inhibitory for RH5 binding, and have used these to understand the molecular basis for the P113:RH5 interaction and to assess the importance of this interaction as a vaccine target.

## Results

### Selection and characterisation of monoclonal antibodies that bind the N-terminal region of P113

To functionally and structurally investigate the interaction between P113 and RH5, we first selected mouse monoclonal antibodies to *P. falciparum* P113. Antibodies were first tested by ELISA to identify those which bind to the N-terminal cysteine-rich region of P113 which contains the RH5 binding site (18), and then for their ability to block binding to RH5. For this study, we selected two anti-P113 monoclonal antibodies: one that could block the interaction with RH5 (10C8), and one that could not (P3.2) (Figure 1). We used surface plasmon resonance (SPR) to quantify the binding of both antibodies to the soluble N-terminal region of P113. Both bound to P113 with nanomolar affinity (10C8, *K*_D_ = 80nM; P3.2, *K*_D_ = 320nM) (Figure 1A, B). Using the AVEXIS binding assay (21), we assessed their ability to prevent P113 from binding to RH5. We found that 10C8 blocked the interaction with RH5, while P3.2 did not (Figure 1C). Because antibodies to RH5 prevent the invasion of erythrocytes by *P. falciparum*, we next asked if there was a correlation between the ability of the anti-P113 antibody to block the RH5 interaction and ability to prevent invasion. By adding serial dilutions of both monoclonal antibodies to a parasite growth inhibition activity (GIA) assay, we found that neither anti-P113 monoclonal antibody was able to inhibit parasite growth in blood culture, even at concentrations that far exceeded the concentrations needed to inhibit the interaction *in vitro*, and at which a monoclonal antibody targeting RH5 (R5.016) (7) showed >90% growth inhibition (Figure 1D).

**Figure 1.**
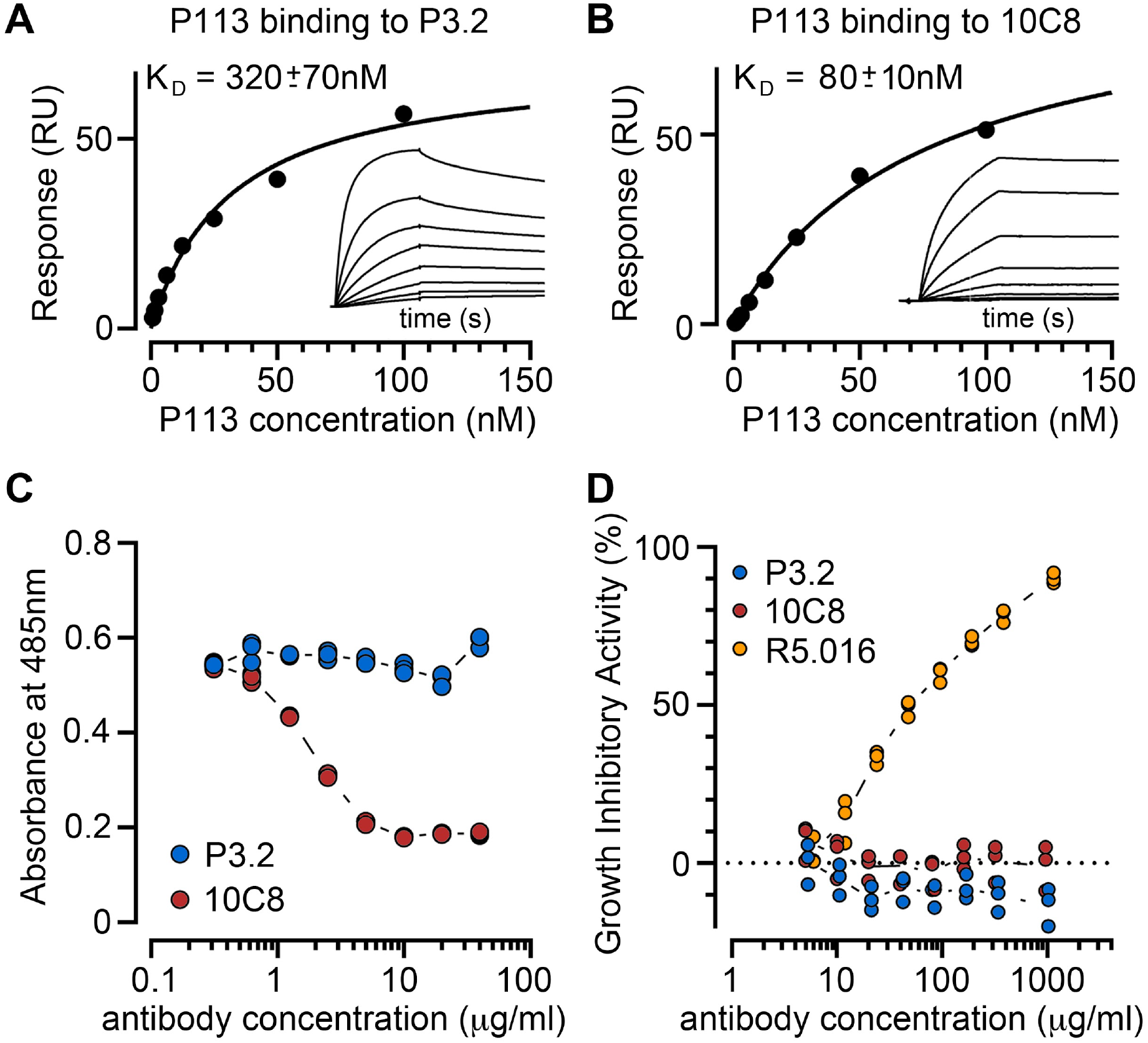
Monoclonal antibodies against P113 that block and do not block RH5 binding do not inhibit parasite growth *in vitro*. Binding kinetics of the anti-P113 monoclonal antibodies P3.2 (**A**), and 10C8 (**B**), to the P113 N-terminal fragment were assessed by surface plasmon resonance. Each monoclonal antibody was immobilised on the sensor surface and the binding parameters of a dilution series of the purified P113 N-terminal fragment was quantified. The response once equilibrium had been reached was plotted and equilibrium dissociation constants (*K*_D_s) were calculated by non-linear curve fitting to the data. Raw sensorgrams are shown inset. **C.** Anti-P113 mAb 10C8 inhibits P113 from binding to RH5 in an AVEXIS assay, but P3.2 does not. The indicated concentrations of protein G-purified monoclonal antibodies were incubated with the biotinylated P113 N-terminal fragment immobilised in wells of a streptavidin-coated microtitre plate before presenting the RH5 beta-lactamase-tagged prey protein; prey binding was quantified by the hydrolysis of the colourimetric beta-lactamase substrate at 485nm. At the antibody concentrations tested, P3.2 did not inhibit the P113-RH5 interaction while 10C8 blocked the interaction. Triplicate data points for each antibody concentration from representative experiments are shown. **D.** Neither RH5 blocking (10C8) or non-blocking (P3.2) anti-P113 monoclonal antibodies inhibit invasion of erythrocytes in a *P. falciparum* blood-stage growth inhibition assay. Synchronised mid-stage trophozoites were added to erythrocytes in the presence of dilution series of 10C8 and P3.2 antibodies. The anti-RH5 mAb R5.016 is included as a positive control. Triplicate data points for each antibody concentration are shown.

### Crystal structure of the N-terminal domains of P113

To better understand the function of the N-terminal region of P113, we determined its crystal structure. A protein containing residues 1-197 was expressed in HEK293 cells and purified for crystallisation. While this did not crystallise alone, we were able to use a Fab fragment of the non-inhibitory P3.2 antibody as a crystallisation chaperone. A complex of P113 1-197 bound to the P3.2 Fab fragment formed crystals that diffracted to 1.95Å resolution. The structure was determined by molecular replacement, using the structure of the Fab fragment of antibody 9AD4 (6) as a search model, followed by iterative cycles of building and refinement, starting with a poly-alanine model of P113. The non-inhibitory antibody, P3.2, bound to an epitope on the narrower side of P113 (Figure 2). Both heavy and light chains are involved in binding, and the epitope has a surface area of ~850Å^2^ (638Å^2^ from V_H_ and 212Å^2^ from V_L_), with the interaction involving fifteen hydrogen bonds, together with surface charge complementarity.

**Figure 2.**
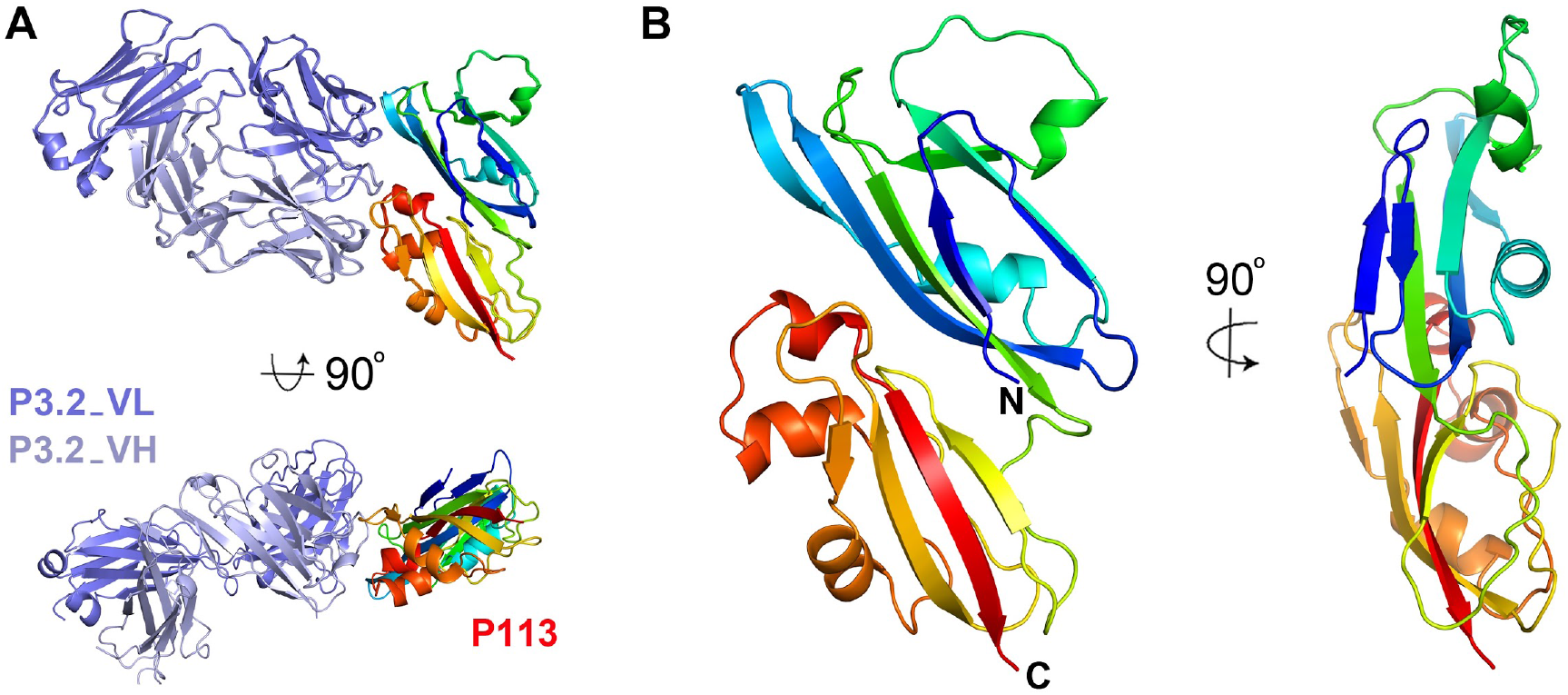
The structure of the N-terminal region of P113. **A.** The structure of residues 1-197 of P113 bound to the Fab fragment of monoclonal antibody P3.2. P113 is shown in rainbow representation, from blue at the N-terminus through to red at the C-terminus. The P3.2 Fab fragment is shown in dark and light blue for the light and heavy chains respectively. **B.** Two views of P113_1-197_, coloured as A.

The structure of the N-terminal region of P113 reveals two closely interacting domains. Both are formed from a four-stranded antiparallel β-sheet which packs against an ɑ-helix (Figure 2, Figure 3). Long loops at one end adopt different structures containing one or two short ɑ-helices with the domains showing an overall root mean square deviation of 3.2Å when compared with each other. In architecture, both domains closely resemble the rhamnose-binding lectin domains, found in proteins such as CSL3 (22), Latrophilin (23) and FLRT (24) (Figure 3). However, structural differences in the region of the rhamnose binding site make it very unlikely that P113 is a rhamnose-binding lectin. Structures of lectin domains from CSL3 (22) and Latrophilin (23), in complex with rhamnose, show that these domains share a GTY motif that orders the loop surrounding the lectin binding site (Figure 3). The GTY motif is lacking in both domains of P113, causing this region of the domain to adopt a different architecture, which does not form a binding pocket for rhamnose. We therefore conclude that despite their architectural similarity, P113 is unlikely to act as a lectin.

**Figure 3.**
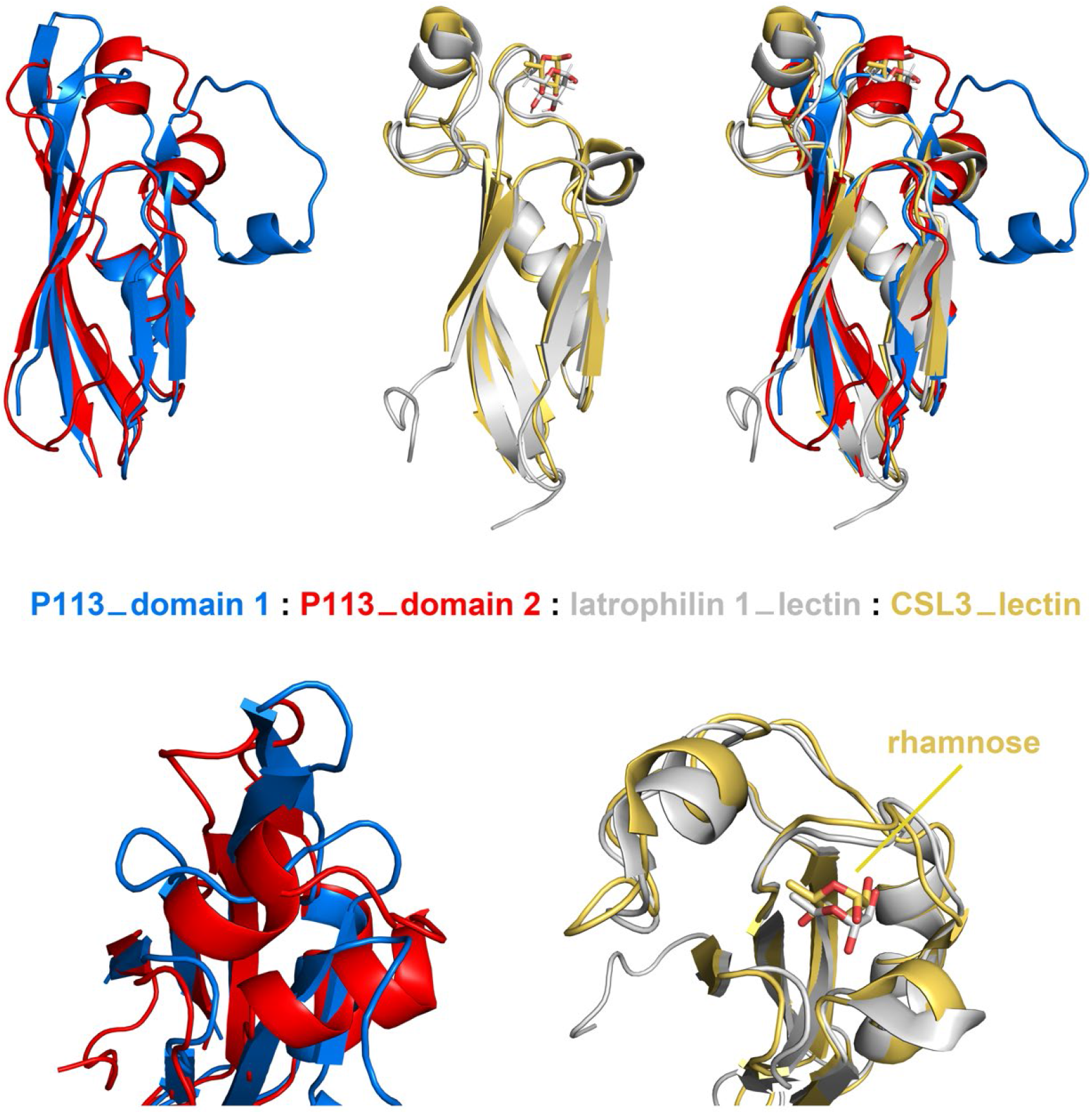
The two domains of P113 resemble rhamnose-binding lectins but lack residues required for rhamnose-binding. A comparison of the structures of domains 1 (blue) and 2 (red) of P113 with rhamnose-binding lectin domains. The structures of the lectin domains of Latrophilin 1 (PDB:2JXA: grey) and CSL3 (PDB: 2ZX2, yellow) are shown in complex with the monosaccharide rhamnose. The lower panel shows a close-up view of the rhamnose-binding pocket of the lectins (right) and the equivalent region of the P113 domains (left).

### Identification of the RH5 binding site on P113

We next aimed to understand the locations of the binding sites on P113 for RH5 and for the inhibitory antibody 10C8. We attempted to crystallise P113_1-197_ in complex with the Fab fragment of 10C8, with RH5, or with a peptide containing residues 9-27 of RH5 previously shown to contain the P113 binding site (18). As these extensive attempts were unsuccessful, we next turned to hydrogen-deuterium exchange mass spectrometry (HDX-MS) to quantify the rate of deuterium exchange of peptides from P113 in the presence and absence of RH5 or 10C8.

The equilibrium binding affinity of full-length RH5 to P113 in solution is 0.3 μM (18); however, addition of RH5 at a concentration of 20 μM caused no significant change in the deuterium exchange of P113 peptides. This lack of protection is consistent with a small binding interface in which the RH5 peptide does not cover a sufficiently large surface area on P113 to alter deuterium exchange. By contrast, we observed reduced deuterium exchange in a number of P113 peptides from residues 103-114 (Figure 4A), including 104-109 (Figure 4B) in the presence of the blocking 10C8 antibody Fab fragment. Labelling these protected peptides on the crystal structure revealed the core of the 10C8 epitope on the surface of P113, which was located on the opposite side of the domain from the non-blocking P3.2 binding site (Figure 4C).

**Figure 4.**
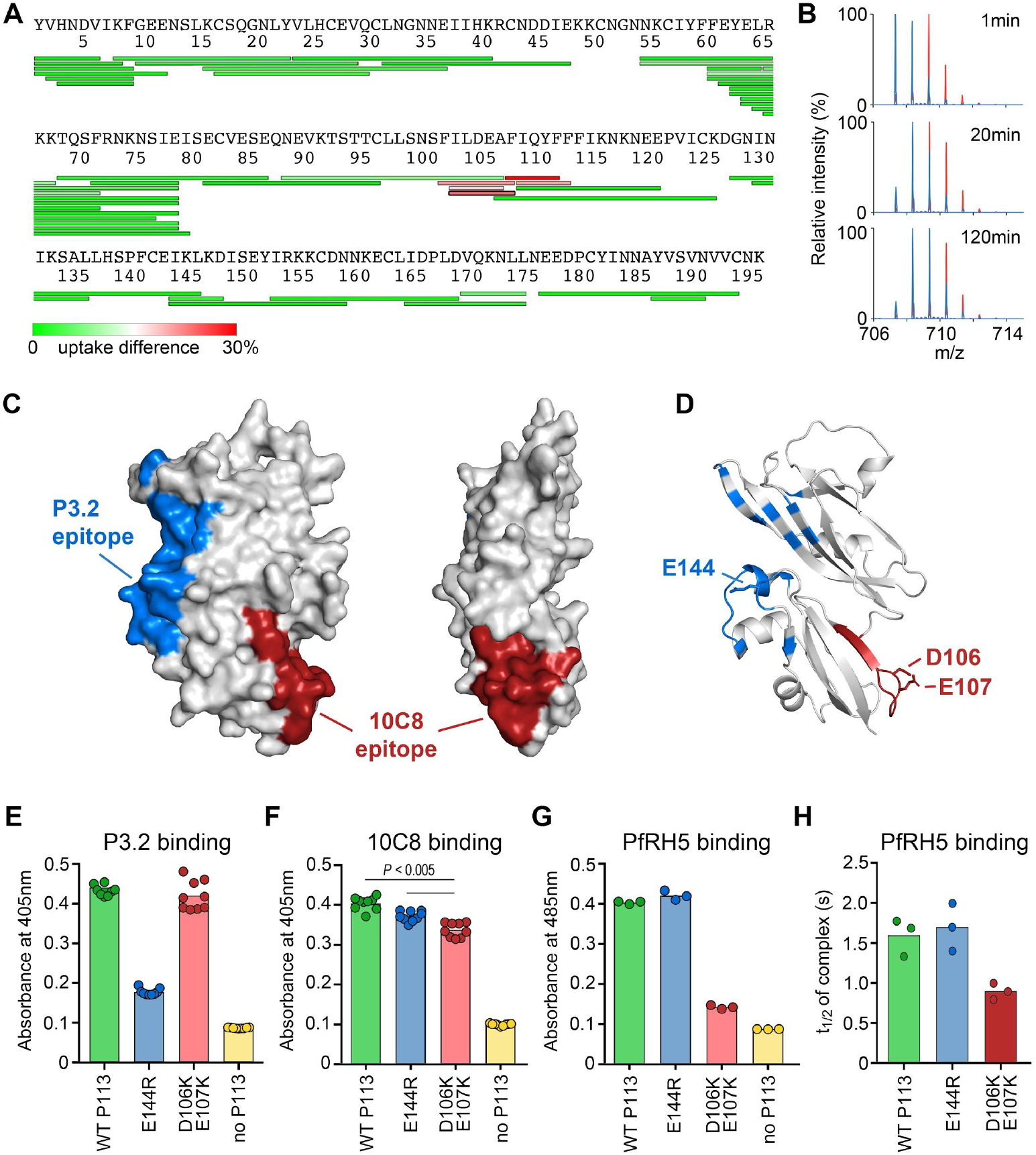
Localisation of the RH5 binding site on P113. **A.** The differences in deuterium uptake of peptides from P113 in the presence of 10C8 after 20 seconds of deuteration. **B.** Mass spectra of P113 peptide 104-109, at different time periods after the start of deuteration in the absence (red) and presence (blue) of 10C8. **C.** A surface representation of the P113 N-terminal domain showing the locations of the anti-P113 mAb epitopes. The epitope for P3.2 as determined from the co-crystal structure, is indicated in blue, and the core of the epitope for 10C8, as determined by HDX-MS mapping, is labelled in red. **D.** The location of mutated residues on the structure of P113. **E, F** Mapping the epitopes of P3.2 (**E**) and 10C8 (**F**) using mutant P113 proteins by ELISA. The indicated mutant and wild-type (WT) P113 N-terminal domains were expressed as biotinylated proteins, immobilised on a streptavidin-coated plate, and the binding of P3.2 and 10C8 quantified by ELISA. The E144R mutant bound 10C8 but not P3.2; the D106K;E107K mutant bound both P3.2 and 10C8, although binding to 10C8 was reduced (unpaired two-tailed t-test, *P* < 0.005). **G.** Location of the RH5 binding site on P113. Biotinylated wild-type and mutant P113 proteins were immobilised on a streptavidin-coated plate and probed for interactions with a pentameric beta-lactamase-tagged RH5 prey protein using the AVEXIS assay. RH5 bound the E144R mutant indistinguishably from wild-type but binding to the D106K E107K mutant was much reduced. **H.** Reduction in binding half-life of RH5 to the D106K E107K mutant as determined by surface plasmon resonance. Interaction half-lives were calculated from the dissociation rate constants determined by fitting the binding data from a dilution series of purified RH5 to the P113 variants to a simple 1:1 binding model. Individual data points from representative experiments are shown; bars represent means.

As antibody 10C8 reduces RH5 binding, we hypothesised that their binding sites overlap. We therefore designed two structure-guided mutants and used the AVEXIS binding assay (21) to test their effect on binding to antibodies P3.2 and 10C8, as well as to RH5. To disrupt 10C8 binding, we selected two neighbouring acidic residues on P113 (at positions 106 and 107), located in a solvent exposed part of the epitope, and mutated both to lysine (D106K E107K). As a control, we made a second mutant with a charge switch mutation within the P3.2 binding epitope (E144R) (Figure 4D). Both mutant proteins were expressed and their ability to bind to P3.2 and 10C8 was determined. The E144R mutation abolished P3.2 binding, confirming the location of the P3.2 epitope (Figure 4E) whereas the D106K E107K mutant caused a small but reproducible reduction of 10C8 binding, demonstrating that the 10C8 epitope was partially affected by this mutation (Figure 4F). Both P113 mutants bound to at least one of the antibodies, suggesting no major disruption to P113 folding.

We then used these mutants to investigate the location of the RH5 binding site on P113 using the AVEXIS assay. We observed that RH5 binding to E144R was indistinguishable from wild-type whereas the binding to the D106K E107K mutant was reduced almost to the level of the negative control (Figure 4G). To confirm and quantify these findings, we measured the binding affinity of a purified monomeric RH5 protein to each mutant using surface plasmon resonance and observed an approximately two-fold reduction in dissociation half-life (Figure 4H). Together, these data localise the binding interface of RH5 on P113; one possible location is the groove formed at the interface of domains 1 and 2, which lies in close proximity to residues D106 and E107 (Figure 4D).

## Discussion

In this study, we analysed two monoclonal antibodies which target P113, allowing us to characterise the molecular basis for the interaction between P113 and RH5. First, we used a Fab fragment from the non-inhibitory antibody, P3.2 as a crystallisation chaperone to determine the structure of the RH5-binding region of P113. This revealed that the N-terminal RH5-binding region of P113 consists of two closely-packed domains, both of which have structural similarities to rhamnose-binding lectins. Despite this architectural similarity, the differences in the domain structure in the location of the rhamnose-binding site make it unlikely that P113 shares their sugar-binding properties.

Despite extensive attempts, we were unable to crystallise P113_1-197_ in complex with either inhibitory antibody 10C8, the P113-binding peptide from RH5, or full-length RH5. Nevertheless, a combination of hydrogen-deuterium exchange mass spectrometry and site-directed mutagenesis allowed us to map the core of their overlapping binding sites on the opposite face of P113 from the P3.2 binding site. A groove immediately adjacent to this 10C8 epitope is a potential binding site for the N-terminal region of RH5.

Our studies also have consequences for the development of vaccines to prevent blood-stage malaria. RH5 and its binding partners have been identified as the most promising candidates for the development of such a vaccine (25, 26). Indeed, RH5 vaccination or passive transfer of RH5 antibodies can elicit protection against *P. falciparum* challenge in non-human primates (10, 11) and vaccination of human volunteers induces production of antibodies with potent growth inhibitory activity (7, 12). As the RH5-binding partners, CyRPA and RIPR are also the targets of antibodies that are inhibitory of parasite growth, this raised the question of whether it is desirable to include P113 as part of a blood-stage malaria vaccine. Supporting this strategy was data which showed that polyclonal antibodies that target the flexible N-terminus of RH5 both prevent P113 binding, and inhibit parasite growth *in vitro* (18).

However, our studies caution against this approach, as a monoclonal antibody which targets P113 and blocks RH5 binding was not inhibitory of invasion of *P. falciparum* into erythrocytes.

How can it be that anti-P113 antibodies which prevent the RH5:P113 interaction are not growth inhibitory? One possibility might be that this interaction is established before P113 is exposed to antibodies on the merozoite surface. Prior to invasion, the proteins of the RH5-containing complex are segregated, with RH5 localising to the rhoptries (17), both CyRPA and RIPR to the micronemes (14), and P113 is embedded in the merozoite plasma membrane (18). It is therefore not known whether these components will be accessible to antibodies before they have formed a stable complex. Indeed, epitopes for structurally characterised RH5 antibodies with growth-inhibitory activity are all found on surfaces of RH5 which remain exposed in its complex with CyRPA and RIPR (6, 7, 19, 25), and lie in the region of the basigin binding site. One possibility is that antibodies that would prevent formation of the RH5:P113 complex cannot access P113 until the complex has already formed. Whatever the reasons, the data presented in this study provides greater insight into the molecular basis for the RH5:P113 interaction but does not support the inclusion of P113 in a subunit blood-stage malaria vaccine.

## Materials and Methods

### Production of protein for binding studies

The N-terminal region of the 3D7 strain *P. falciparum* P113 protein was expressed by transient transfection of HEK293 cells using the expression plasmids described in (20). Briefly, the P113 expression plasmids were chemically synthesized using codons optimised for expression in human cells, with potential N-linked glycosylation sites mutated, and with a C-terminal rat Cd4d3+4 tag (27). Where appropriate, monomeric “bait” proteins were enzymatically monobiotinylated by co-transfection with a plasmid encoding a secreted BirA enzyme (21, 28). To make mutations in the P113 sequence, PCR primers were designed with the intended nucleotide change and site directed mutagenesis performed using KOD Hot Start DNA polymerase, as per the manufacturer’s instructions. His-tagged proteins were purified from supernatants on HisTrap HP columns using an ÄKTApure (GE Healthcare) and resolved by gel filtration on a Superdex 200 Increase 10/300 column (GE Healthcare) (29).

### Protein production for crystallisation

To produce P113_1-197_ for crystallisation studies, the gene fragment comprising the first 591bp of the *P113* gene sequence, previously codon optimised for expression in *H. sapiens* (GeneArt), was sub-cloned into the P113-bio vector (Plasmid 47729, Addgene) using a Gibson Assembly Cloning Kit (NEB). Primers were designed to introduce a thrombin site between the *P113* gene fragment and the coding sequence for Cd4-His_6_ on the P113-bio vector. The resulting plasmid DNA construct was used to transfect Expi293F human cells using the Expi293F transfection kit and Expi293F expression medium (Thermo Fisher). Cells were harvested by centrifugation at 1,000 g for 30 min and resuspended in 1 x PBS supplemented with 30 mM imidazole. Cell lysis was performed using a cell disruptor at 10 kpsi pressure and the cell lysate was centrigued at 50,000 g for 30 minutes at 4 °C. The soluble fraction was incubated for 1 hour at 4 °C using a Ni-NTA resin (Qiagen). The Ni-NTA resin was washed with 4 times resin volume (RV) of Buffer A followed by elution with 3 RV of 1 x PBS supplemented with 0.5 M imidazole. The eluted protein was desalted against 1 x PBS buffer and digested with thrombin protease (GE Healthcare) in a ratio of 1 unit per μg at room temperature. The reaction mixture was loaded into a gravity flow column pre-loaded with Ni-NTA beads to remove Cd4-His6 obtaining tagless P113_1-197_.

### Monoclonal antibody generation

To generate monoclonal antibody P3.2, female BALB/c mice were used (Harlan Laboratories, Oxfordshire, UK). All procedures on mice were performed in accordance with the terms of the UK Animals (Scientific Procedures) Act Project Licence and were approved by the University of Oxford Animal Welfare and Ethical Review Body. Mice were immunised intramuscularly (IM) with 20 μg of the entire ectodomain of P113 (18) adjuvanted in a 1:1 ratio of Addavax (Invivogen, cat no. vac-adx-10) followed by two similar IM boosts at 2 week intervals, followed by a final intraperitoneal (IP) boost in PBS, two weeks later. Three days after the final IP boost, the mice were culled and spleens and blood collected. Splenocytes were isolated by shredding the spleen through a mesh followed by washing three times in ClonaCell-HY Medium B (StemCell technologies). Splenocytes were fused to SP2/0 cells using the ClonaCell-HY hybridoma Kit (StemCell cat. 03800). In brief, washed spleen cells and SP2/0 cells were mixed in a 5:1 ratio, pelleted, and carefully resuspended in ClonaCell-HY PEG solution followed by centrifugation 133 x g for 3 min. PEG solution was aspirated and cells resuspended initially in ClonaCell-HY Medium B thereafter in ClonaCell-HY Medium C (StemCell technologies) and incubated overnight (37°C, 5% CO_2_). On the following day, cells were pelleted (10 min 400 x g), resuspended in 10 mL ClonaCell-HY Medium C and 90 mL ClonaCell-HY medium D (semisolid HAT-selection media), and plated out into 100 mm petri plates. On day 13, colonies were picked manually and transferred to HT-selection media. Culture supernatants were screened for specificity by ELISA and positive cultures were single cell sorted using a Beckman Coulter Legacy MoFlo MLS High Speed Cell Sorter (BD). Monoclonal cultures were expanded in DMEM (Sigma) containing L-glutamine, Pen/strep and ultra low FCS, and antibodies were purified from the supernatant by gravity flow protein G columns. For monoclonal antibody production, hybridoma cells were pelleted at 1000 x g and the supernatant containing antibody P3.2 was filtered with 0.22 μm filter units (Nalgene) before being loaded onto Protein G resin (Thermo Fisher) previously equilibrated with 20 mM HEPES, 150 mM NaCl pH 7.4. The resin was subsequently washed with 3 resin volumes (RV) of the same buffer before mAb P3.2 (Thermo Fisher) was eluted using 3 RV of 0.2 M glycine pH 2.5. The pH of the solution was immediately neutralised with 1M Tris-HCl pH 7.5 to a final concentration of 50 mM and mAb P3.2 was desalted in 1 x PBS. Papain digestion was performed overnight at RT using the Pierce Mouse IgG1 Fab and F(ab’)2 Kit (Pierce) and Fc was separated from Fab fragments using protein A resin. Fab P3.2 containing aliquots were stored at −80 °C in 1 x PBS. Monoclonal antibody 10C8 was prepared in a similar way except SJL-strain mice were co-immunized with both full-length P113EE and P113-Y1-N653 proteins (18). Immunized mice were tested for P113-specific antibody titers by ELISAs essentially as described (30). Hybridoma supernatants were screened for binding to P113EE, P113-Y1-N653 and P113-Y1-K197 proteins by ELISA. Further, clones were counter-screened against the rat Cd4-His protein to remove any antibodies reacting against the tag. The 10C8 monoclonal antibody was purified on a Protein A column using standard methods. The antibody was eluted at pH 3.0 and dialysed against PBS. The dialyzed antibody was sterile filtered, isotyped as IgG1/kappa. Binding to P113EE and P113-Y1-N653 protein was verified by ELISA.

### Enzyme-linked immunosorbent assay (ELISA)

Protein expression levels and monoclonal antibody binding were quantified by ELISA essentially as described (31). Briefly, biotinylated proteins were immobilised in individual wells of streptavidin-coated microtitre plates and the appropriate primary antibody added. To quantify protein expression levels, the mouse anti-rat Cd4 monoclonal antibody (OX68) which recognises the Cd4 expression tag was used, and to determine the location of antibody epitopes, the anti-P113 mAbs P3.2 and 10C8 were added. An anti-mouse-alkaline phosphatase conjugate was used as a secondary antibody (Sigma, UK).

### AVidity-based EXtracellular Interaction Screening (AVEXIS)

AVEXIS screening was performed essentially as described (21). Briefly, monomeric biotinylated bait proteins and highly avid pentameric β-lactamase-tagged prey protein were prepared and their expression levels normalised using enzyme activity to approximately 5μg mL^−1^ prior to their use in interaction screening (28). Biotinylated baits were immobilised in streptavidin-coated 96-well microtitre plates, washed with PBST (PBS/0.1 % Tween-20), incubated with prey proteins, and washed three times with PBST. Captured preys were quantified by adding the colorimetric β-lactamase substrate nitrocefin and measuring the absorbance of the hydrolysis products at 485 nm. The negative control in each screen was the query prey protein probed against the Cd4d3+4 tag alone.

### Surface plasmon resonance analysis

Surface plasmon resonance studies were performed using a Biacore 8K instrument (GE Healthcare) as described (20). Biotinylated bait proteins were captured on a streptavidin-coated sensor chip (GE Healthcare). Approximately 400 RU of the negative control bait (biotinylated rat Cd4d3+4) were immobilised on the reference flow cell and approximate molar equivalents of the query protein immobilized in other flow cells. Purified analyte proteins were separated by size exclusion chromatography on a Superdex 200 Increase 10/300 column (GE Healthcare) in HBS-EP just prior to use in SPR experiments to remove any protein aggregates that might influence kinetic measurements. Increasing concentrations of purified proteins were injected at 100 μL/min to determine kinetic parameters, or at 30 μL/min for equilibrium measurements. Both kinetic and equilibrium binding data were analysed in Biacore 8K evaluation software version 1.1 (GE Healthcare). Equilibrium binding measurements were taken once equilibrium had been reached using reference-subtracted sensorgrams. Both the kinetic and equilibrium binding were replicated using independent protein preparations of both ligand and analyte proteins. All experiments were performed at 37°C in HBS-EP (10 mM HEPES, 150 mM NaCl, 3 mM EDTA, 0.05% v/v P20 surfactant).

### Determination of the structure of P113 bound to monoclonal antibody 3.2

P113_1-197_ was combined with Fab P3.2 in a 1:1.2 w/w ratio for 1 h at room temperature before lysine methylation method as described (32). Residual methylation reagents were neutralised by addition of 1M Tris-HCl pH 7.5 to a final concentration of 50 mM Tris-HCl and the protein complex was concentrated to 2 mg mL^−1^ using 30 kDa cut-off concentrators (Millipore) before injection into a S75 16/600 column (GE Healthcare) previously equilibrated with 20 mM Tris-HCl, 150 mM NaCl, pH 7.4. Fractions containing the P113_1-197_:Fab P3.2 complex were pooled and concentrated to 10 mg mL^−1^ as appropriate for crystal formation based on use of pre-crystallisation kit (Hampton Research). Crystallisation trials in 96-well plates were set-up at 4 °C with the sitting-drop method using commercial screens (ProPlex, Midas, Morpheus, MembFac and MembGold2) with 100 nL reservoir + 100 nL protein drops. Diamond-shaped crystals appeared in 7-10 days in 0.1 M Mg acetate, 0.1 MES pH 5.8, 22 % w/v PEG4000 and were supplemented with 20% w/v MPD before being flash cooled in liquid N_2._ Two datasets from a single crystal were collected at 100K at beamline I03 of the Diamond Light Source. The crystal belonged to space group P4_1_2_1_2 with unit cell dimensions a=97Å,b=97Å, c=178Å and α=β=γ=90.00°. The datasets were indexed and reduced to 1.95 Å resolution using autoPROC (33). The R_free_ set, comprising 5% of the reflections, was generated randomly in Unique. The structure was solved by molecular replacement using as searching models the light and heavy chain of Fab 9AD4 (PDB ID: 4U0R), after pruning the side chains using chainsaw (CCP4 package, (34)) and manually deleting the variable regions in the PDB. The two searching models were used sequentially in PHASER as implemented in Phenix MR (35). The MR solution was further extended using BUSTER (36) using the missing atoms option leading with initial R_factor_ and R_free_ of 41 and 45% respectively.

The majority of P113_1-197_ (163/197aa) was initially built as poly-alanine model in PHENIX Autobuild, followed by multiple runs of restrained refinement in PHENIX (resolution range 50.58-1.95 Å). Reiterated model building was performed in COOT (37) and structure validation was performed using MOLPROBITY (38) before deposition in the PDB (PDB accession code: 6Z2L). Crystallography data and refinement statistics are reported in Table 1. All the crystallographic programmes were used as part of the SBGrid package (39).

**Table 1:** crystallographic statistics.

|  | **P113<sub>1-197</sub>-Fab P3.2** |
| --- | --- |
| **Beamline** | Diamond I03 |
| **Wavelength (Å)** | 0.9762 |
| **Resolution range (Å)** | 63.95 - 1.95 (2.02 - 1.95) |
| **Space group** | *P* 4<sub>1</sub> 2<sub>1</sub> 2 |
| **Unit cell parameters** | a= 96.93Å b= 96.92Å c=177.90Å,<br>α=β=γ= 90.00° |
| **Total reflections** | 124960 (12324) |
| **Unique reflections** | 62481 (6162) |
| **Multiplicity** | 2.0 (2.0) |
| **Completeness (%)** | 99.8 (100.00) |
| **Mean I/sigma(I)** | 17.31 (1.80) |
| **Wilson B-factor (Å<sup>2</sup>)** | 41.80 |
| **R-merge** | 0.0162 (0.439) |
| **R-meas** | 0.0229 (0.620) |
| **R-pim** | 0.0162 (0.439) |
| **CC1/2** | 1 (0.655) |
| **CC\*** | 1 (0.890) |
| **Reflections used in refinement** | 62477 (6162) |
| **Reflections used for R-free** | 3109 (300) |
| **R-work** | 0.204 (0.302) |
| **R-free** | 0.225 (0.310) |
| **CC(work)** | 0.944 (0.735) |
| **CC(free)** | 0.945 (0.684) |
| **Number of non-hydrogen atoms** | 5329 |
| **macromolecules** | 4929 |
| **ligands** | 5 |
| **solvent** | 395 |
| **Protein residues** | 631 |
| **RMS bonds (Å)** | 0.012 |
| **RMS angles (°)** | 1.35 |
| **Ramachandran favored (%)** | 97.56 |
| **Ramachandran allowed (%)** | 2.44 |
| **Ramachandran outliers (%)** | 0 |
| **Rotamer outliers (%)** | 0.71 |
| **Clashscore** | 3.5 |
| **Average B-factor (Å<sup>2</sup>)** | 43.14 |
| **macromolecules** | 42.53 |
| **ligands** | 79.98 |
| **solvent** | 50.3 |
| **PDB ID** | 6Z2L |

Structure homology was analysed using DALI server (40), whilst protein-protein interface was analysed using PISA (41). Protein Topology was produced using PDBsum (42) and protein images were produced using the graphic software PYMOL (43).

### Hydrogen-deuterium exchange mass spectrometry

HDX-MS was performed using a Waters HDX manager composed of a nano-Acquity UPLC coupled to a Synapt G2-Si (Waters) mass spectrometer. Samples were prepared by 11-fold dilutions from 7 μM of apo P113 in deuterated or non-deuterated and P113-10C8 or P113-RH5 complex in deuterated 20 mM HEPES, 150 mM NaCl pH 7.4 buffer. The pH of the sample was reduced to 2.3 by adding 50% vol/vol 150 mM HCl. The apo protein and complexes were digested in-line using a pepsin-immobilised column at 20 °C. The peptides generated from pepsin digestion were trapped on a micro peptide trap for 2 min at a flow rate of 200 μL min^−1^, to allow the removal of salts and were then separated using a C18 column with a linear gradient of 5-65% acetonitrile in water, both supplemented with 0.1 % formic acid, for 9 min at flow rate of 40 μL min^−1^. The temperature at which liquid chromatography temperature was performed was set at 0 °C to reduce back-exchange. Peptide mapping of P113 was performed by using non-deuterated samples in triplicate and only unique peptides present in all three data files were selected for deuterium uptake data analysis. Apo P113 protein digestion provided a list of 1,649 peptides. After applying selection filters, and after manual inspection, 59 peptides were selected for analysis. These peptides provided ~95% sequence coverage with many overlapping peptides. For labelling experiments, apo P113, P113-10C8 or P113-RH5 were incubated for 20 s, 10 min and 2 h in deuterated buffer. All HDX-MS experiments were performed in duplicate. Sequence coverage and deuterium uptake were analysed by using ProteinLynx Global Server (Waters) and DynamX (Waters) programmes, respectively. Leucine enkephalin at a continuous flow rate of 5 μL min −1 was sprayed as a lock mass for mass correction.

### *P. falciparum* culture and growth inhibition activity assays

The ability of anti-P113 mAbs to inhibit *in vitro* growth of *P. falciparum* 3D7 parasites was assessed using a standardized procedure. The mAbs were buffer-exchanged against incomplete culture medium (25 mM Hepes, 0.37 mM hypoxanthine, and 2 mM L-glutamine in RPMI (Sigma, R0883)), concentrated with centrifugal filter devices (Amicon, Fisher Scientific, UFC901096) and sterilized by filtration through a 0.22 μm spin filter (Costar Spin-X, SLS Ltd, 8160) before use in the assay. These samples were tested in a serial 3-fold dilution with a start concentration of 1 mg mL^−1^ IgG in triplicate in a one-cycle growth inhibition assay (GIA) using human O+ erythrocytes parasitized with mid-trophozoite stages of *P. falciparum* prepared by 5% sorbitol treatment on the previous day. Parasite growth after approximately 44 h of culture was determined by a biochemical assay specific for parasite lactate dehydrogenase (44). Values obtained with test IgGs were compared with those obtained with parasites incubated in the presence of positive and negative controls (normal growth medium, 5 mM EDTA, positive (RH5-specific) and negative control mAbs).

## Acknowledgements

This research was funded by Wellcome Trust (grant 206194) (GJW, FG) and a Wellcome Investigator Award (101020/Z/13/Z) to MKH. SJD is a Jenner Investigator, Lister Institute Research Prize Fellow and a Wellcome Trust Senior Fellow (106917/Z/15/Z). SM and APS are supported by the Francis Crick Institute, which receives its core funding from Cancer Research UK (FC001999), the UK Medical Research Council (FC001999), and the Wellcome Trust (FC001999). The funders had no role in study design, data collection and interpretation, or the decision to submit the work for publication. TWP and VK are employees of Leidos, Inc., the prime contractor for the Malaria Vaccine Development Program (MVDP), Contract AID-OAA-C-15-00071, with the Office of Infectious Diseases, Bureau for Global Health, U.S. Agency for International Development (USAID), which provided funding for development of the 10C8 mAb. The opinions expressed herein are those of the authors and do not necessarily reflect the views of the U.S. Agency for International Development.

## Author contributions

IC solved the crystal structure of the P113-Fab complex; FG performed the protein and antibody binding assays. SM and KEW performed the HDX analysis under the supervision of APS. LKB cloned the P3.2 antibody, DQ performed the GIA assays, and both were supervised by SJD. TWP and VK provided the 10C8 mAb and information regarding its development and characterization. MKH and GJW managed the project and analysed data. IC, FG, MKH and GJW wrote the manuscript.

## Data availability

The coordinates and structure factors associated with this work are available at the Protein Data Bank (PDB: 6Z2L). All other data are available from the authors on request.

